# Genetic and transcriptomic analysis of hyphal constriction based on a novel assay method in the rice blast fungus

**DOI:** 10.1101/2023.06.14.545034

**Authors:** Eunbyeol Cho, Song Hee Lee, Minsoo Jeong, Surajit De Mandal, Sook-Young Park, Seung Won Nam, Do Gyeung Byeun, Jung Kyu Choi, Yong-Hwan Lee, Jae-Ho Shin, Junhyun Jeon

## Abstract

An ability of fungi to undergo hyphal constriction is important for fungal ecology and diseases. In the rice blast fungus, *Magnaporthe oryzae*, hyphal constriction is required to traverse host cell junctions through pit fields. However, little is known about genetic underpinnings of hyphal constriction, except the requirement of a mitogen-activated protein kinase, Pmk1. Here we demonstrate that a simple *in vitro* assay based on nitrocellulose membrane allows investigation of the genetic basis for hyphal constriction. Using the assay, we found that the constriction limit of *M. oryzae* hypha lies between 0.22 and 0.3 μm, and that a histone modification might be involved in hyphal constriction. RNA-seq experiments combining our assay and Δ*pmk1* showed that hyphal constriction relies on transcriptional changes of genes implicated primarily in membrane and cell wall-related cellular processes in Pmk1-dependent and/or Pmk1-independent manner. Furthermore, our assays with diverse fungal species suggest correlation between hyphal constriction and fungal lifestyles. Our study reveals that hyphal constriction can be induced without host-derived cues and provides molecular and evolutionary insights into a fundamental process to infection of host plant.

## Introduction

An ability to undergo hyphal constriction is vital for fungi to explore the confining environments, penetrate into host cells, and traverse cell junctions^1,2^. Hyphal constriction is a cell morphological process of filamentous fungi that might be parallel to the interstitial migration of mammalian cells, in that both processes require severe deformation of cell and its contents^2-4^. Despite its apparent implication in fungal biology and evolution of eukaryotic traits, genetic basis of hyphal constriction remains largely unknown. Such scarcity of our knowledge on hyphal constriction is due in large part to the difficulty of monitoring hyphal constriction in natural settings such as soil and host tissue. Although a few studies investigated the behavior and plasticity of fungal hyphae using fabricated microenvironments, none of them focused on hyphal constriction ^1,5,6^.

*Magnaporthe oryzae* (synonym: *Pyricularia oryzae*) is a filamentous fungal pathogen that causes the rice blast disease. *M. oryzae* also infects other major cereal crops including wheat^7^. Given that the rice blast destroys enough rice to feed more than 60 million people and puts the world’s wheat yield at risk, the fungus poses a serious threat to global food security^8^. Infection of host plants by *M. oryzae* usually begins with germination of a disseminated 3-celled asexual spore, conidium. During germination, the germ tube tip differentiates into a specialized infection cell, appressorium within 4 to 6 hours. The appressorium develops enormous turgor pressure of up to 8 MPa, which is channeled as mechanical force into a narrow penetration peg in order to breach the cuticular layer of leaf surface^7,9^. Once inside a plant cell, the penetration peg differentiates into bulbous invasive hyphae that are encased in plant plasma-membrane^10,11^. At this stage, the fungus rapidly fills the interior of the first host cell and spread to the adjacent cells through pit fields where plasmodesmata cluster, resulting in colonization of host tissue^10,12,13^.

During the infection and colonization of host plants by *M. oryzae*, there are two processes that require the constriction of hyphae: penetration peg formation and cell-to-cell movement^2^. To date, extensive work has been dedicated to uncovering and characterizing many genes that regulate appressorium formation, as this morphogenetic process is the first critical step for invasion into the host cells^14^. Decades of works have revealed that signaling pathways implicating cAMP, calcium, and MAP kinases are pivotal for differentiation of appressorium^14-16^. Recently, it was demonstrated based on elegant combination of pharmacological and genetic tools that Pmk1 (MAP kinase regulating appressorium development and pathogenicity), not Mps1 (cell wall integrity MAP kinase), is required for *M. oryzae* hyphae to undergo severe constriction (from an average diameter of 5 μm to 0.6 μm) in order to transverse from one cell to the neighboring cells^17^. As Mst12, which is involved in penetration peg formation, is the transcription factor phosphorylated by Pmk1, this attests to the importance of the MAP kinase pathway for hyphal constriction^18^. However, genetic underpinnings of hyphal constriction are largely unknown. Most of the genetic mutations in signaling pathways and other cell regulatory genes preclude the appressorium differentiation, rendering it difficult to examine the contribution of the genes to the later processes.

Here we show that a simple assay based on the nitrocellulose membrane provides a genetic screen method for hyphal constriction. Using the assay system in combination with a variety of mutant strains and RNA-seq experiment, we investigated genetic underpinning of hyphal constriction in *M. oryzae*. We also probed the possible correlation between hyphal constriction and fungal lifestyles. The resulting insight into this important yet understudied fungal development would be relevant to not only better understanding of fungal pathogenicity and its evolution but also agricultural and medical applications.

## Results

### Limitation of microfluidic chips in screening hyphal constriction

In previous studies, microfluidic chips were used to investigate the growth pattern and foraging behavior of fungal hyphae, as they provide both flexibility in creating different environments and feasibility in tracking the hyphae under the microscope^5,6^. In order to test whether we can take advantage of the microfluidic platform for monitoring of hyphal constriction, we attempted to compare the wild-type strain with *Pmk1* and *Mck1* (an upstream MAP kinase of Mps1) null mutant strains on the microfluidic chips fabricated to have a narrow passage point with diameter of approximately 2 μm (Supplementary Fig. 1). It was technically, if not impossible, extremely challenging to fabricate micro-channels having much less than 2 μm in diameter. When the fungal hyphae on the chips were monitored under the microscope following inoculation of different *M. oryzae* strains at the one end of the chips filled with the complete media, we found that they were able to traverse the narrow passage point without much difficulty, regardless of the strains (Supplementary Fig. 1). This result suggests that the 2 μm channel is not narrow enough to reveal the differences in ability to undergo extreme hyphal constriction.

### Development of the nitrocellulose membrane-based assay system

Due to the technical limitation of the microfluidic chip, we turned to the nitrocellulose membrane (NM), which is easily accessible and comes in a variety of pore sizes starting from 0.22 μm. We hypothesized that the different pore sizes of the NM might be able to differentiate the mutant strains for extreme hyphal constriction. We set out to probe this possibility by devising a simple assay, in which an ability to undergo hyphal constriction can be visually examined (Fig. 1A). The key idea is that if the fungus is capable of traversing the membrane of a particular pore size, then the fungal colony would form on the underlying medium right beneath the point of passage after the removal of NM, providing a visual readout for successful passage.

Prior to comparing the wild-type and the mutant strains using our assay, we first tested the passage of the NM by a wild-type strain, KJ201 over a range of pore sizes by incubating an agar block on top of the membrane for two days (Supplementary Fig. 2). This showed that the KJ201 is able to pass through the membrane of 0.33 μm and larger pore, but not the membrane of 0.22 μm. When we tested different wild-type strains, however, there seemed to be variation in the readouts: CP987 and KI215 were comparable to the KJ201, while 70-15 and Guy11 were unable to form colonies beneath the 0.45 μm membrane (Supplementary Fig. 3A). When the test was performed again, accounting for the differences in hyphal growth rate among the wild-type strains (Supplementary Fig. 3B) by incubating the agar blocks for three days instead of two, 70-15 and Guy11 formed colonies as well (Supplementary Fig. 3C). This result indicated that incubation period should be adjusted for comparison between wild-type and mutant strains, depending on which wild-type strain is used and whether or not the mutant has growth defect.

**Figure 1.**
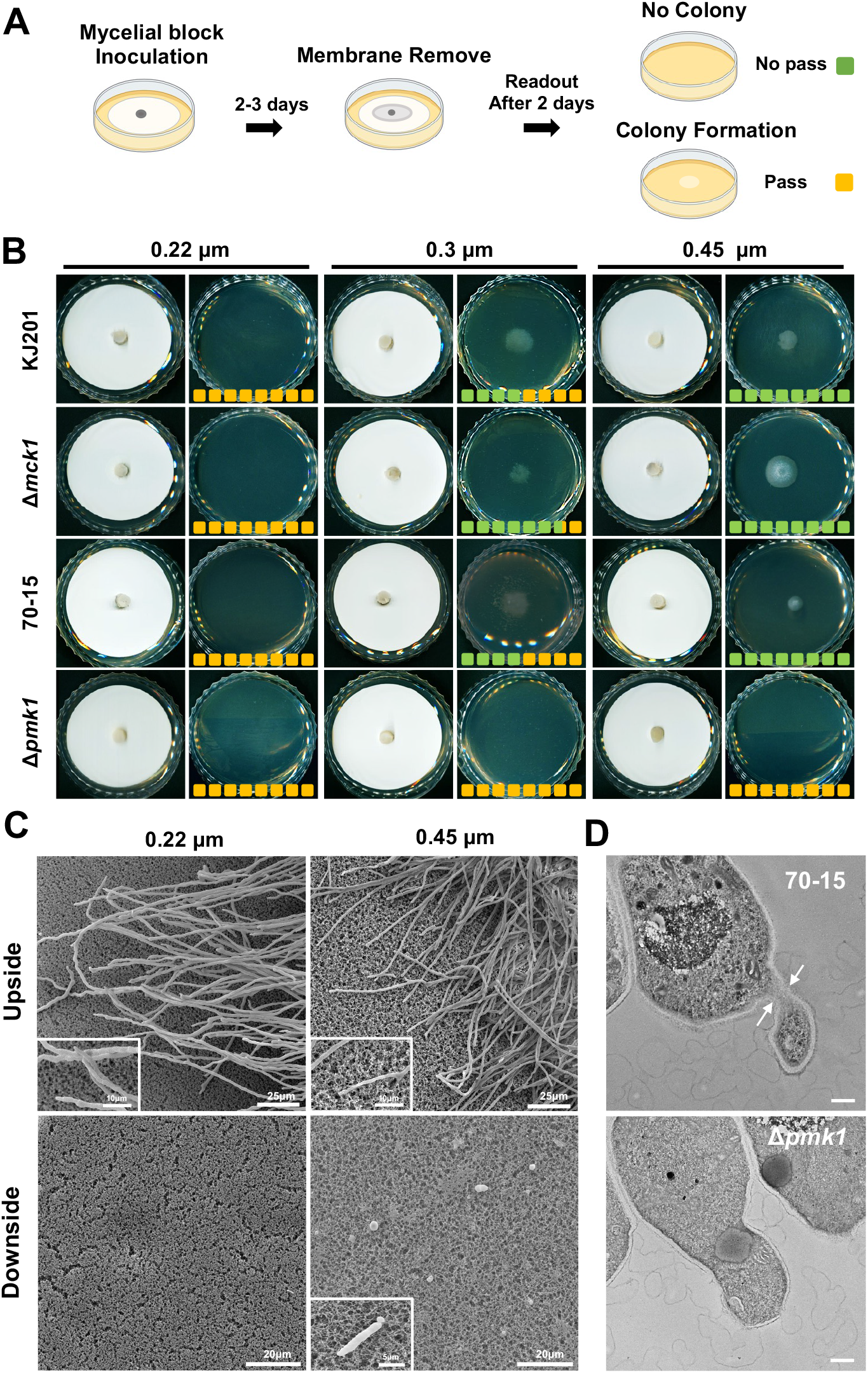
Hyphal constriction can be induced and tested using nitrocellulose membrane-based assay. **A**, In nitrocellulose membrane (NM)-based assay, agar plug (grey circle) is placed upon the membrane on the complete medium agar plate. Following 2-3 days of incubation depending on the growth rate of strains tested, the NM is removed from the plate, which is incubated for two more days. Passage of NM by fungal hyphae can be judged visually by colony formation on the medium. Green and orange boxes represent the color codes used in B for the two visual readout (no passage and passage, respectively). **B**, Using the NM of varying pore sizes, different strains of *M. oryzae* including Δ*pmk1*, which is defective in hyphal constriction, were tested for their ability to pass through the NM. The number of boxes indicates the number of replications, and colors represent assay readouts. **C**, Scanning electron microscopy shows upside and downside of 0.22 and 0.45 μm membranes during the NM-based assay. A few hyphae coming out of the membrane surface are observed on the downside of 0.45 μm NM, but none are visible on 0.22 μm NM. Insets show magnified view of designated area. **D**, Transmission electron microscopy reveals hyphal constriction of 70-15 strain inside the matrix of 0.45 μm NM, in contrast to the Δ*pmk1* showing no hyphal constriction.

Next, we asked if our assay is able to differentiate the wild-types and mutants in their ability to transverse the membrane (either 70-15 and Δ*pmk1* pair or KJ201 and Δ*mck1* pair) (Fig. 1B). Clearly, all the strains could not pass through the 0.22 μm membrane. For 0.3 μm membrane, KJ201 showed mixed results, while 70-15 didn’t seem to pass through it at all. Such observation suggests that there might be subtle but inherent differences in ability to undergo hyphal constriction among wild-type strains, and that limit of hyphal constriction lies between 0.22 and 0.3 μm. In our assay, the Δ*pmk1* did not show any sign of membrane passage even for 0.45 μm membrane, unlike Δ*mck1* that was comparable to KJ201 in membrane passage. This result is consistent with the reported role of *Pmk1* in hyphal constriction.

### Validation of the assay system as a tool for testing hyphal constriction

As our assay is based on the assumption that the fungus forms a colony only when it can pass through the membrane by undergoing hyphal constriction, we first checked the possibility that the fungus makes its way through the membrane via action of cellulases encoded in the genome. Making a mutant lacking cellulase activity was not feasible, since *M. oryzae* has multiple copies of cellulase genes. Moreover, silencing those genes is very likely to confound the interpretation of the assay outcomes by a significant amount of residual cellulase activity. Instead of the genetic approach, therefore, we tested if the fungus is able to pass through nylon membrane or polyvinylidene difluoride (PVDF) membrane, which also comes in a variety of pore sizes. The assay using such extremely hard-to-degrade membranes (0.45 μm) showed that the fungus can pass through those membranes as well (Supplementary Fig. 4), suggesting that enzymatic break-down of materials contribute little, if at all, to the membrane passage.

In order to ascertain that the fungus passes through the membrane via hyphal constriction, we probed the membrane via both scanning and transmission electron microscopy (SEM and TEM). Our SEM analysis on the membrane surfaces showed that there are no appressoria on upside of the membrane, and that hyphae emerge out from downside of 0.45 μm membrane but not from that of 0.22 μm (Fig. 1C). The SEM image also showed that the membranes do not provide regular grid structure but matrix of irregular size of pores. Nevertheless, TEM analysis on the longitudinal section of membranes revealed that 70-15 strain could undergo the extreme hyphal constriction down to approximately 0.5 μm (Fig. 1D upper panel), while in stark contrast, Δ*pmk1* could only go into the membrane without hyphal constriction at a site where the pore size is large enough to accommodate the hyphae (Fig. 1D lower panel). These observations indicate that 1) hyphal constriction can be induced without host cell-derived cues and 2) our NM-based assay allows to distinguish the abilities of the different strains to undergo hyphal constriction.

### Screening of mutant lines impaired in signaling pathways and histone modifications

Our data demonstrating that the fungus passes through the membrane when it is capable of hyphal constriction prompted us to screen mutant lines of *M. oryzae* using our NM-based assay (0.45 μm pore membrane). We have screened two categories of mutant lines available to us (signaling pathway mutants and histone modifying enzyme mutants) in an attempt to understand how the fungus genetically control hyphal constriction. Our screening of signaling pathway mutants indicated that Pmk1 appeared to be the primary regulator of hyphal constriction (Fig. 2A and Supplementary Fig. 5). Although cAMP signaling pathway is absolutely required for appressorium formation, none of the related mutants seemed to be defective in membrane passage.

**Figure 2.**
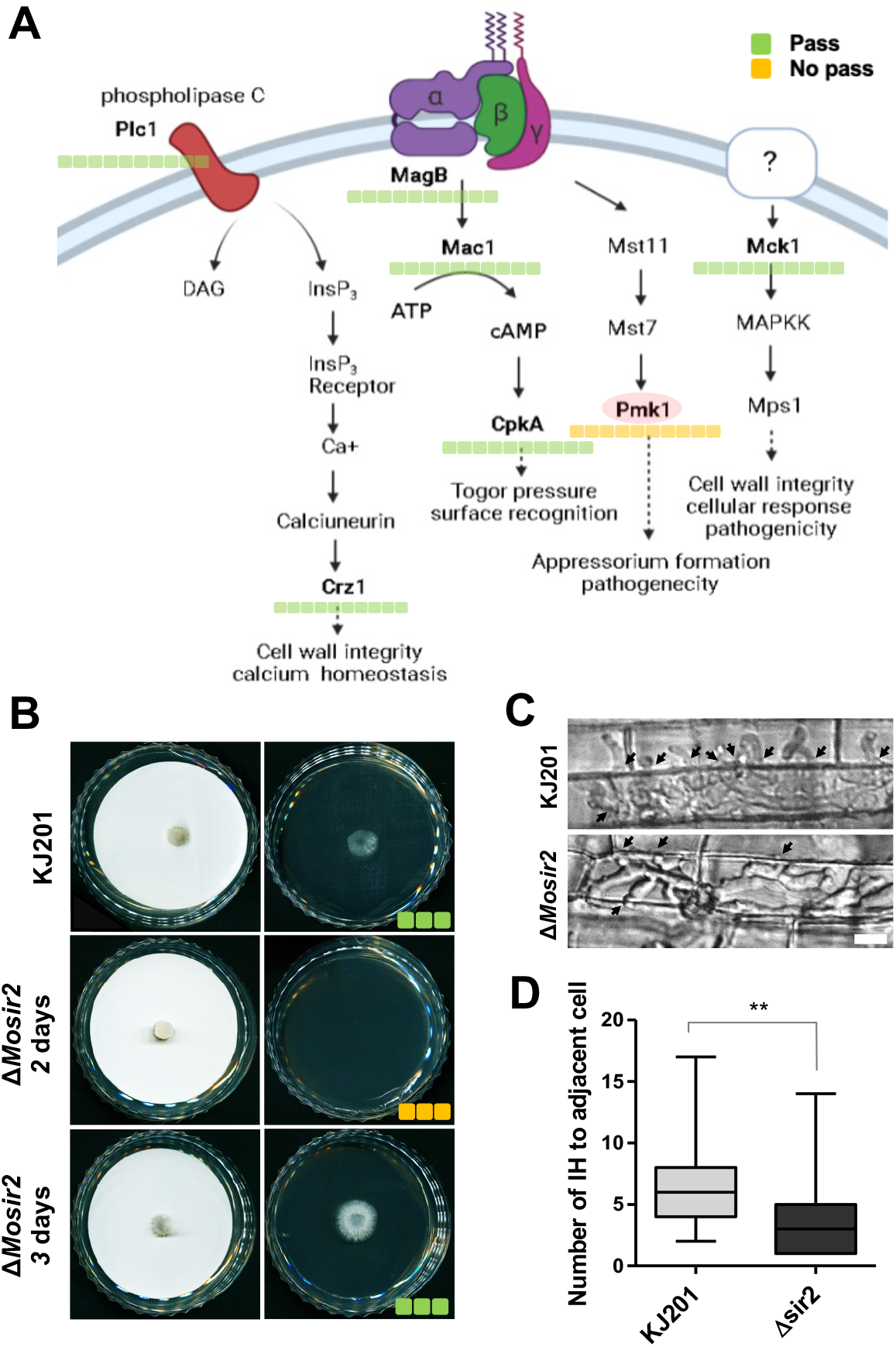
NM-based assay establishes *Pmk1* as a central and *MoSir2* as an auxiliary regulator of hyphal constriction in the rice blast fungus. **A**, Diagram showing genes and components of different signaling pathways and NM-based assay readouts for the mutants of selected signaling pathway genes. Pmk1 is shaded in pink, as it is the only gene of which deletion completely blocks the membrane passage. **B**, NM-based assay readouts for KJ201 and Δ*Mosir2*. The Δ*Mosir2* passed through the NM in 3-day incubation setting, but it did not in 2-day setting, despite the comparable growth rate as the wild-type. **C**, Sheath assay was performed with KJ201 and Δ*Mosir2*, and the number of cell-to-cell movement of each strain to the adjacent cells was counted at the first invaded cells. **D**, A boxplot shows the distribution of the number of invasive hyphae (IH) moving to adjacent cells (Wilcoxon rank sum test, *P* < 0.01).

Screens of histone modifying enzyme mutants showed that most of genes we tested are dispensable for hyphal constriction, except the gene encoding a class III histone deacetylase, *Sir2* (Sirtuin-2) (Fig. 2B and Supplementary Fig. 6). The Δ*sir2* was reported to be comparable to the wild-type strain in hyphal growth and appressorium function, but it was shown to be defective in suppressing plant defense response and colonization due to the failure to regulate superoxide dismutase gene expression^19^. Since the Δ*sir2* seemed to be inefficient in membrane passage in our assay, we tested the cell-to-cell movement Δ*sir2* using rice sheath cells treated with NADPH oxidase inhibitor diphenyleneiodonium (DPI) which allows evaluation of cell-to-cell movement of the mutant by suppressing the reactive oxygen burst as previously described^19^ (Fig. 2C). In the experiment, the Δ*sir2* showed significantly less numbers of invasive hyphae that traverse cell-cell junctions than the wild-type, suggesting that it is indeed compromised in hyphal constriction (Fig. 2D). Overall, our screens of mutant lines in *M. oryzae* indicate that the NM-based assay can be a powerful tool in isolating and characterizing the genes involved in hyphal constriction and cell-to-cell movement of fungal pathogens.

### Transcriptome analysis reveals genomic basis of hyphal constriction

To investigate how hyphal constriction is regulated at the genomic level, we designed and performed the RNA-seq analysis, taking advantage of our assay system and Δ*pmk1* in combination (Fig. 3A). RNAs were extracted either from hyphae scraped off the complete media agar plates or from hyphae contained within the nitrocellulose membranes (see the Materials and Methods for details). Gene expressions were compared among samples using an adjusted *P*-value < 0.05 and fold change of two as criteria for differential expression. Differential expression of genes in our dataset was validated by qRT-PCR analysis for randomly selected 16 genes (Supplementary Fig. 7). Comparison of 70-15 inside membrane to 70-15 on CM (denoted as c/a) showed 350 (approximately 2.7% of genome) differentially expressed genes (DEGs), of which 167 and 183 genes were up-regulated and down-regulated, respectively (Fig. 3B and Supplementary Data 1). Comparison of 70-15 and Δ*pmk1* in membrane (denoted as d/c) revealed 501 DEGs (226 up-regulated and 275 down-regulated), while comparison of 70-15 and Δ*pmk1* in CM (denoted as b/a) revealed 1,181 DEGs (687 up-regulated and 505 down-regulated genes) (Fig. 3B and Supplementary Data 1).

**Figure 3.**
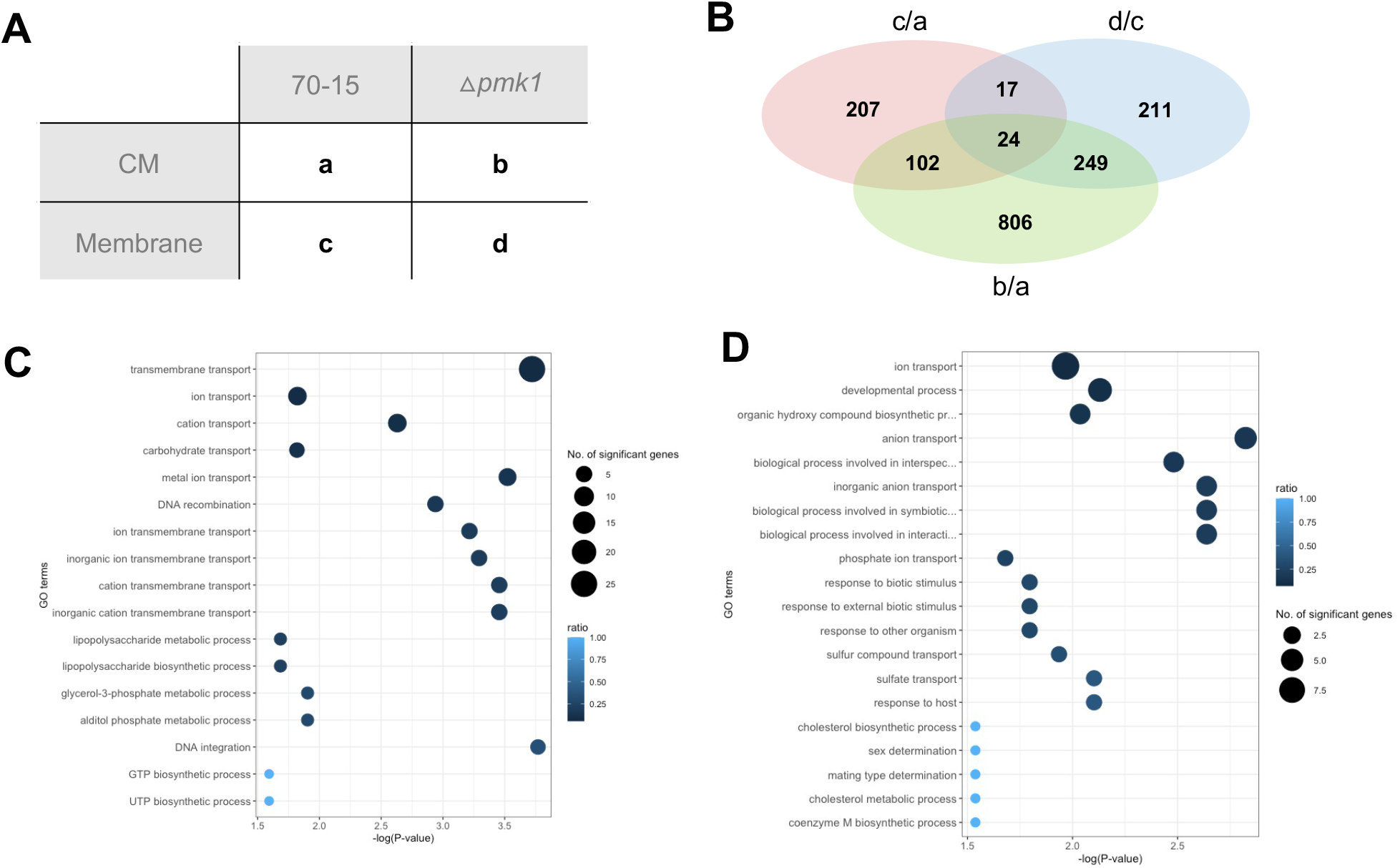
Transcriptome analysis using NM-based assay reveals *Pmk1*-dependent and/or *Pmk1*-independent genes involved in hyphal constriction. **A**, RNA samples were prepared from four different settings combining two strains (70-15 and Δ*Mosir2*) and two culture conditions (CM and membrane). Each setting was labeled as a to d. **B**, Venn diagram shows the number of differentially expressed genes (DEGs) among three comparisons. The c/a represents comparison of c relative to a. The d/c represents comparison of d relative to c. The b/a represents comparison b relative to a. **C and D**, Summary of GO enrichment analysis (biological process) for selected terms. Top 20 GO terms (based on *P*-value) that are enriched with DEGs from c/a and d/c are shown (Fisher’s exact test, *P* < 0.01). The area of the circle represents the number of genes assigned to the particular GO term. The color of the circle indicates the proportion of genes assigned to the GO term in our dataset among the total number of genes having that GO term in the genome.

Recently, Yan *et al*. provided a high-resolution transcriptional profiling of the plant-associated development of the rice blast fungus, in which they detected 10 modules depending on temporal co-expression pattern of genes^20^. Comparison of our DEG sets to genes in individual modules revealed the significant overlaps of our DEGs with modules 6, 8 and 9, all of which correspond to stages of rice blast disease development involving cell-to-cell movement via hyphal constriction (Supplementary Fig. 8). This observation suggests that our membrane-based assay offers *in vitro* condition that mimics *in vivo* cell-to-cell movement via hyphal constriction.

In our transcriptome analysis, c/a, b/a, and d/c were expected to reveal genes involved in hyphal constriction genes, Pmk1-dependent genes in CM, and Pmk1-dependent genes in membrane, respectively. Overlaps between any two sets of DEGs were all statistically significant, pointing to high similarities among transcriptional changes in comparisons of samples (hypergeometric test: *P* = 9.7e-10 between c/a and d/c, *P* = 1.3e-44 between c/a and b/a, and *P* = 4.4e-156 between b/a and d/c). Gene ontology (GO) analysis (biological process) showed that DEGs from c/a are enriched with GO terms associated with transmembrane transport of ions and metals, DNA recombination, membrane synthesis, while over-representation of genes associated with developmental process and interaction with other organisms was a main feature of DEGs from d/c (Fig. 3C, D and Supplementary Fig. 9). The DEGs from b/a showed enrichment of GO terms associated with a wide range of metabolic processes (Supplementary Fig. 9). GO analysis on cellular component indicated the enrichment of DEGs in membrane and cell wall, regardless of comparisons (Supplementary Fig. 10). Overall, our transcriptome analysis suggests that hyphal constriction requires both Pmk1-independent regulation of membrane-related processes/functions and Pmk1-dependent regulation of genes involved in development and interaction with host, along with transcriptional changes of genes implicated in diverse cellular metabolisms. Interestingly, DEGs (24 genes) that are shared by all three combinations (Supplementary Data 2) included the tetrahydroxynaphthalene reductase gene (MGG_02252, *Buf1*)^*21*^.

### Fungal lifestyles correlate with capability of hyphal constriction

Fungi display different lifestyles: saprobes, biotrophs, hemibiotrophs, and necrotrophs. *M. oryzae* is a hemibiotrophic pathogen that can switch from biotrophic to necrotrophic lifestyle. One interesting question is whether the ability to undergo extreme hyphal constriction is an inherent and universal characteristic of fungal species, and whether it is related to lifestyles. In an attempt to gain insight into this question, we tested the membrane passage (0.45 μm pore size) by saprobic, necrotrophic, and hemibiotrophic fungal species. For each species, the incubation period was adjusted according to the measured growth rate. The assays for a total of fourteen species suggested strong association between the lifestyles and the hyphal constriction (Fig. 4 and Supplementary Fig. 11). While all the saprobes and hemibiotrophs were able to pass through the membrane, the necrotrophs rarely made their way through the membrane. To exclude the possibility that membrane passage is dependent on how wide or thick the fungal hyphae are, we measured the width of hyphae for all the species tested, and found that the dimension of hyphae is not a determining factor of the membrane passage. This result suggests that hyphal constriction might be a fungal trait in close association with lifestyles. *Fusarium graminearum*, which has been traditionally considered as a necrotrophic fungal pathogen but was recently classified as a hemibiotroph due to its brief biotrophic phase during infection and colonization, was able to pass through the membrane in our assay^22^. Interestingly, *F. graminearum* was shown to pass between host cells via pit fields by undergoing hyphal constriction^23^. This may suggest that membrane passage trait could be a predictor of fungal lifestyle.

**Figure 4.**
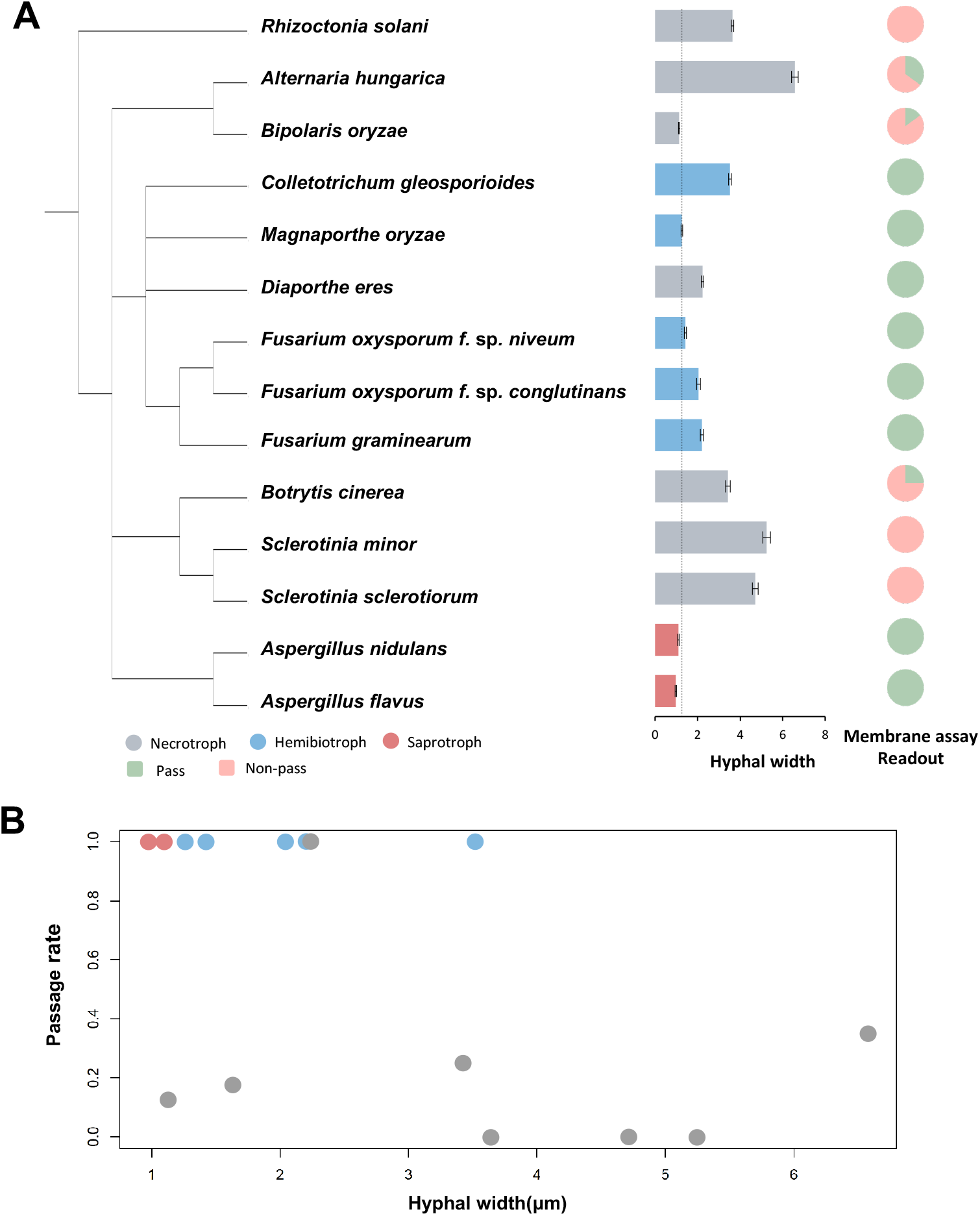
Hyphal constriction correlates with lifestyle of fungi. **A**, fungal species with different lifestyles were ordered by phylogenetic relationship, and measured hyphal widths and membrane assay readouts of individual species were plotted as bar graph and pie charts. Dotted line in the bar graph indicates the hyphal width of *M. oryzae*. **B**, Scatterplot shows the relationship between hyphal width and membrane passage. Lifestyle of fungi were color-coded as in A.

## Conclusion

Taken together, our works demonstrate that nitrocellulose membrane-based assay can be an invaluable tool for studying genetic components underlying the important fungal development. Microscopic and genetic analysis of membrane passage show that hyphal constriction can be induced without an obvious plant-derived molecular cues. The transcriptome analysis taking advantage of our assay method suggest that the membrane passage via hyphal constriction is a complex process that is parallel in many aspects to what happens *in planta*. The nitrocellulose membrane has been used in many of *Agrobacterium*-mediated transformation of fungi for random insertional mutagenesis. Our data point to the limitation of such experiments, as the use of membrane can be discriminate against the mutants that are unable to pass through it. We believe that use of our assay system in *M. oryzae* and other fungi would shed light on a fundamental yet understudied fungal development for pathogenesis.

## Materials and Methods

### Fungal strains and culture conditions

The wild type *M. oryzae* strains used in this work were KJ201,70-15, GUY11, CP987, KI215. The mutant *M. oryzae* strains used in this study were Δ*pmk1* (70-15 background), Δ*mck1* (KJ201 background), Δ*magB* (70-15 background), Δ*cpka* (70-15 background), Δ*plc1* (KJ201 background), Δ*crz1* (KJ201 background), Δ*rpd3* (KJ201 background), Δ*hos2* (KJ201 background), Δ*jmj1* (KJ201 background), Δ*sas3* (KJ201 background), Δ*jmjd2* (KJ201 background), and Δ*sir2* (KJ201 background). The following fungal strains were used to test correlation between hyphal constriction and lifestyles: *Rizoctonia solani, Bipolaris oryzae, Aspergillus flavus, Aspergillus nidulans, Botrytis cinerea, Sclerotinia sclerotiorum, Sclerotinia minor, Colletotrichum gloeosporioides, Fusarium graminearum, Fusarium oxysporum f. sp. conglutinans*, and *Fusarium oxysporum f. sp. niveum. M. oryzae* strains were maintained on complete medium agar (CM, 0.6% yeast extract (w/v), 0.6% casamino acid (w/v), 1% sucrose (w/v) and 1.5% agar (w/v); 25 °C). Other fungal strains were maintained on potato dextrose agar (1.5% agar (w/v), MB CELL, Seoul, Korea). All plate images were taken using a MICROTEK XT3300 scanner at a resolution of 800dpi.

### Microfluidic chip hyphal constriction screening

The 3-channel microfluidic chip was manufactured to test hyphal constriction. The microfluidic chip was designed to be 10 μm in width by autoCAD (Autodesk), and fabricated using PDMS (polydimethylsiloxane 182 base and curing agent, Dowsil, CA, USA) attached to slide glass. The chip has three narrow channels (2 μm) for observation, with two openings (1.5 mm in diameter for each) at either side of slide for inoculation and nutrient supply. The chips were inoculated with the fungal strains and incubated at 25°C for 24 h to observe passage of hyphae through the narrow channels. Imaging was performed using Leica DM2500 microscope and Leica DFC7000T digital camera.

### Nitrocellulose membrane assay

Nitrocellulose membrane (GVS, white disk diam. 47 mm, hydrophilic) was placed upon complete medium agar in 60×15 mm of petri-dish. Cylindrical agar block was placed on the membrane and incubated for 2 to 4 days depending on growth rate of strains tested. Following incubation, the membrane with agar block on it was removed, and plate was incubated 2 days for readout. Assays using nylon membrane (GVS, white disk diam. 47mm, hydrophilic) and PVDF membrane (Durapore, disk diam. 47 mm, hydrophilic) were done following the same protocol as nitrocellulose membrane.

### Electron microscopy

For scanning electron microscopic (SEM) imaging, 70-15 and Δ*pmk1* strain was inoculated upside of nitrocellulose membrane on CM plate and incubated for 3 days as described. The membranes were taken from the plates and then fixed in 5% glutaraldehyde at room temperature for 1 h. The fixed samples were treated with 1% osmium for 1 h and dehydrated in an ethanol series (50%, 60%, 70%, 80%, 90% and 100%) followed by drying using an E3100 Critical Point Dryer (Quorum Technologies Ltd., Laughton, UK). The dried filters were mounted on stubs and coated with platinum. The samples were observed with a Mira-3 FE-SEM (Tescan Ltd., Brno, Czech) at 5–10 kV. For transmission electron microscopy (TEM), samples were prepared as described for SEM sample fixation, and embedded in epoxy. Serial thin sections of embedded sample were placed onto carbon grid and examined with a Hitachi H-7600 transmission electron microscope.

### Rice sheath assay and imaging

Conidia were collected from 9-day-old oatmeal agar cultures and resuspended to 5 × 10^4^ conidia per ml in deionized water supplemented with 0.4 μM DPI (NADPH oxidase inhibitor diphenyleneiodonium, sigma, MA, USA). Rice sheath segments were obtained from four-week-old susceptible rice seedlings (Nakdongbyeo). Spore suspensions were gently injected into 4 ∼5 cm-long rice sheaths and incubated in plates covered with a wet paper towel for high humidity for 25°C for 48 h. Imaging of infected sheaths was performed after 48 hpi, using Leica DM2500 microscope and Leica DFC7000T digital camera.

### RNA-seq sample preparation and analysis of RNA-seq data

For RNA extraction, nitrocellulose membrane assay (0.45 μm) method was modified as follows: instead of agar block, ground mycelia were spread on the membrane and incubated. Following incubation, mycelial masses on the upside of membrane was scraped off, and the membranes containing mycelia inside them were collected and grinded with mortar and pestle in liquid nitrogen. Total RNAs were extracted using Easy-spin total RNA extraction kit (iNtRON Biotechnology, Seoul, Korea). In RNA-seq experiment, paired-end sequencing of mRNAs was carried out on the Illumina platform. The quality of the raw fastq reads derived from the sequencing was checked by FastQC (Andrews S., 2010, https://github.com/s-andrews/FastQC) and processed by TrimGalore (https://github.com/FelixKrueger/TrimGalore) to remove the low-quality bases and adapter sequences. Trimmed paired-end reads from each sample were aligned to the *M. oryzae* reference genome with STAR aligner^24^. Differential expression of genes was analyzed with the DESeq2 Bioconductor package ^25^. The threshold selected for differentially expressed genes (DEGs) was fold-change of 2 and adjusted P-value < 0.05. Gene ontology was carried out using the topGO package ^26^ and enrichment of genes in particular GO terms was analyzed using Fisher ’ s exact test (*P* < 0.05).

## Acknowledgement

This work was supported by grants from the National Research Foundation of Korea (NRF-2018R1A5A1023599 and 2021R1A2C2012002).

