## Supplementary Fig for "Genetic and transcriptomic analysis of hyphal constriction based on a novel assay method in the rice blast fungus"

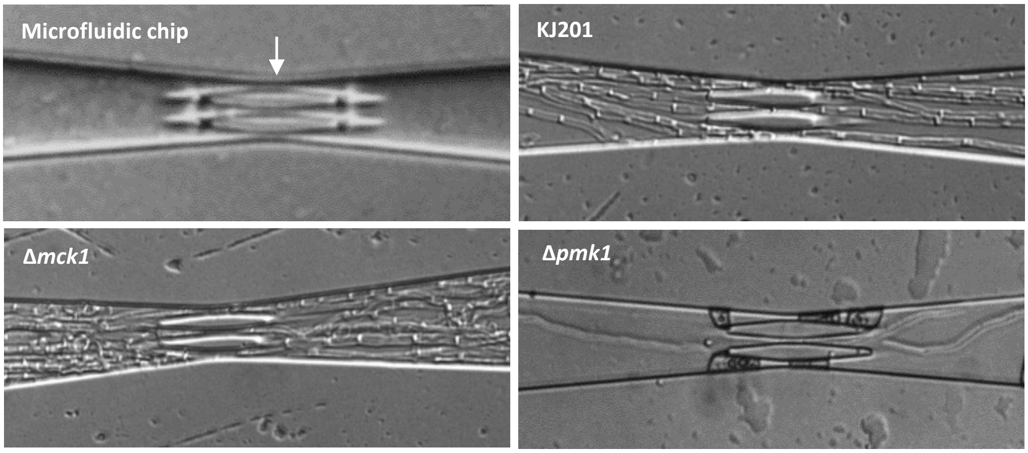


**Supplementary Fig. 1**. Testing microfluidic chip for hyphal constriction. Microfluidic chip was fabricated to have three narrow (2 µm) channels in the middle of the chip (indicated by an arrow), which is filled with complete media. Chip was inoculated with fungi at its one end, and then incubated overnight for observation under the microscope. All the strains tested appeared to pass through the channels.

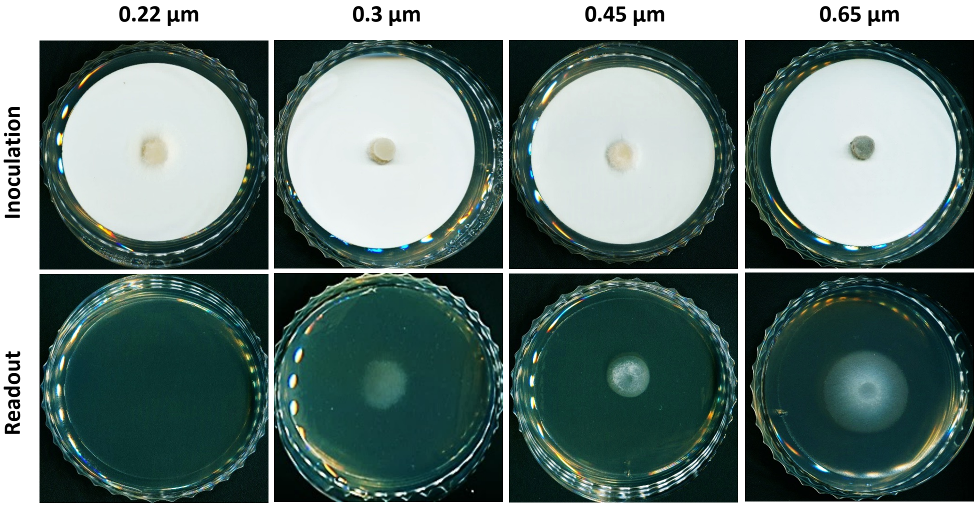


**Supplementary Fig. 2**. Readouts from nitrocellulose membrane-based assay for KJ201 using membranes of varying pore size.

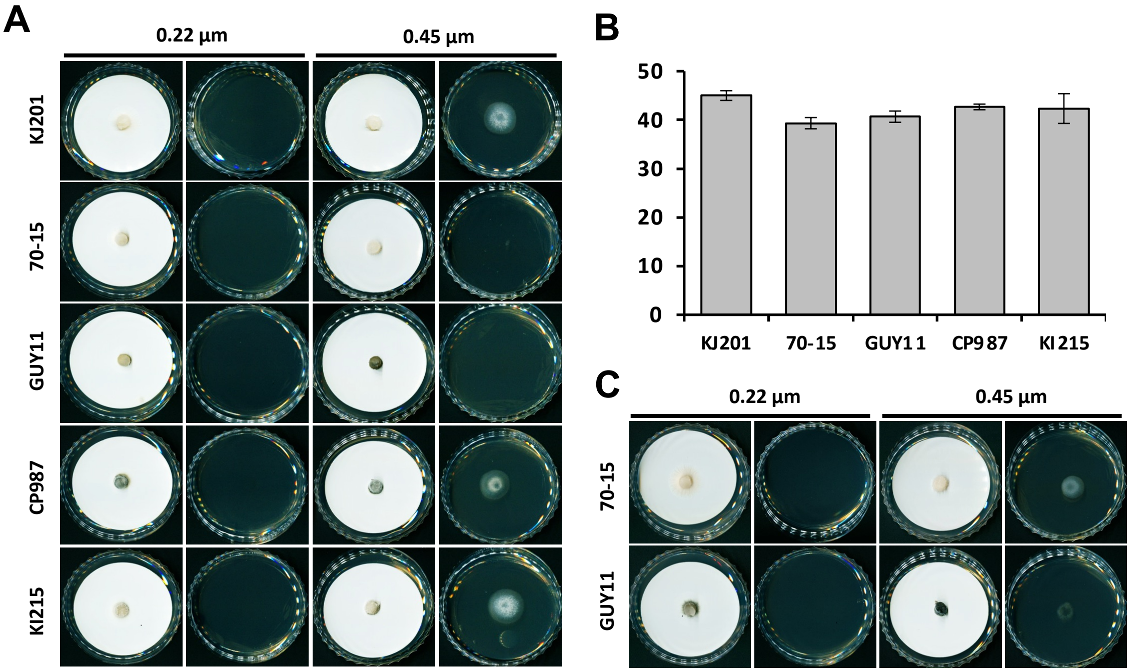


**Supplementary Fig. 3**. Adjustment of assay condition based on the growth rate. **A**, Different wild-type strains shows differences in their ability to pass through the membrane. Agar block was placed on the membrane and incubated for two days before removal of membrane. **B**, Bar graph showing inherent differences in growth rate of wild-type strains. **C**, To account for the differences in growth rate, 3-day instead of 2-day of incubation period was used for 70-15 and Guy11, resulting in positive readout in the assay.

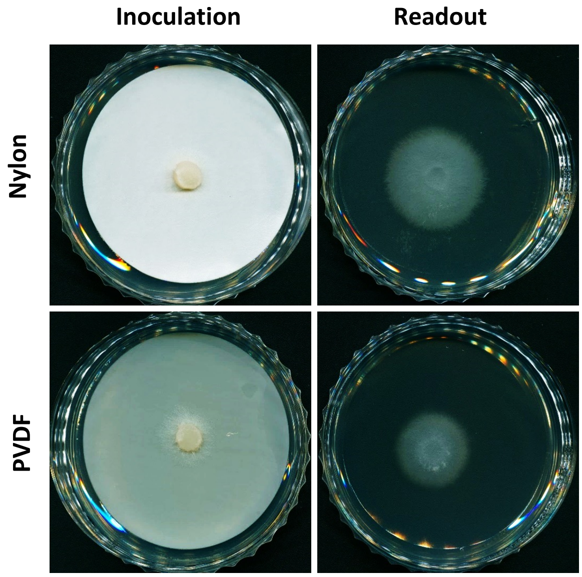


**Supplementary Fig. 4**. Use of other types of membrane for the assay. Assays using nylon or PVDF membrane having pore size of 0.45 µm instead of nitrocellulose membrane resulted in 'positive' readout, suggesting that cellulase activity is not responsible for passage of fungal hyphae through the membrane.

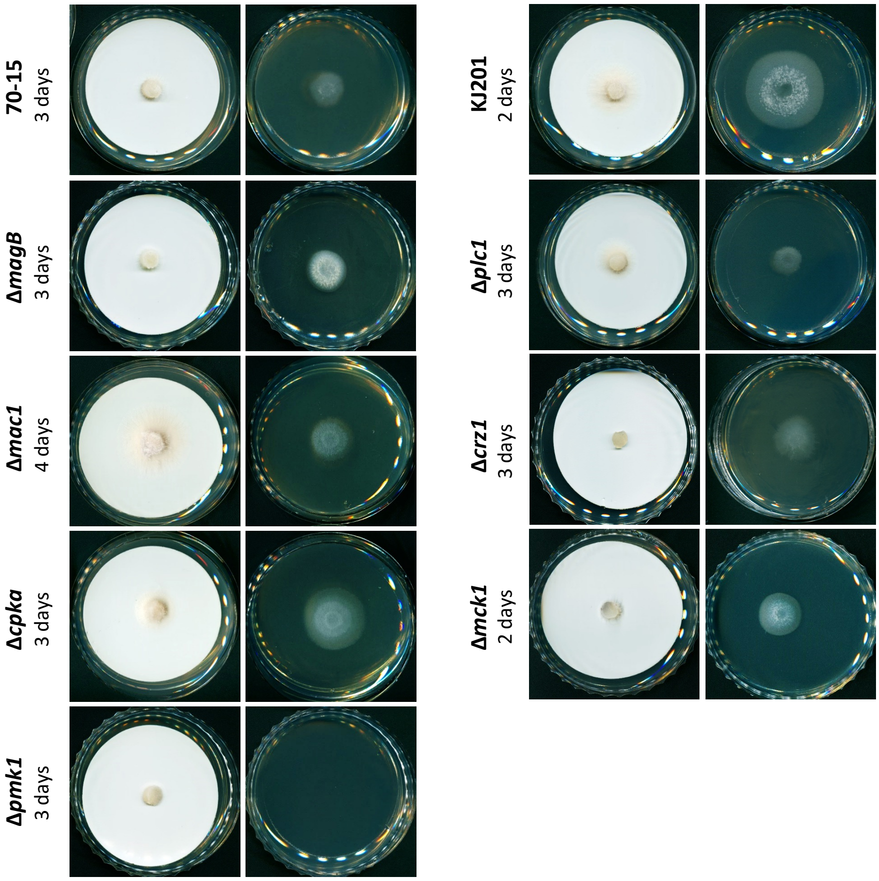


**Supplementary Fig. 5**. Screening of mutants involved in signaling pathways for their ability to pass through the membrane. Initial incubation period was adjusted according to the growth rate. The *Pmk1* deletion mutant was the only one that showed negative readout.

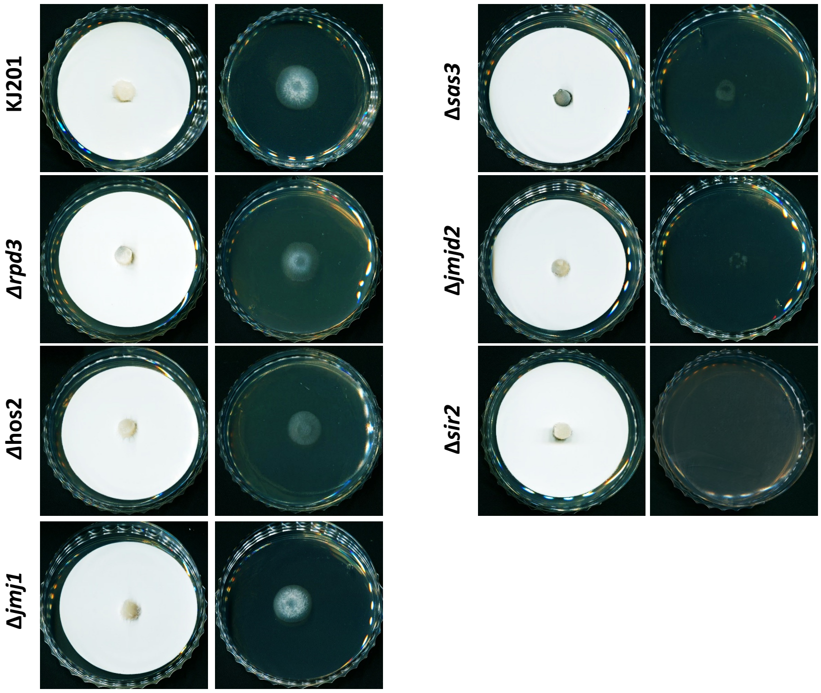


**Supplementary Fig. 6**. Screening of the mutants involved in histone modifications for their ability to pass through the membrane. The *Sir2* deletion mutant showed negative readout in the initial screening.

### RNA-seq validation

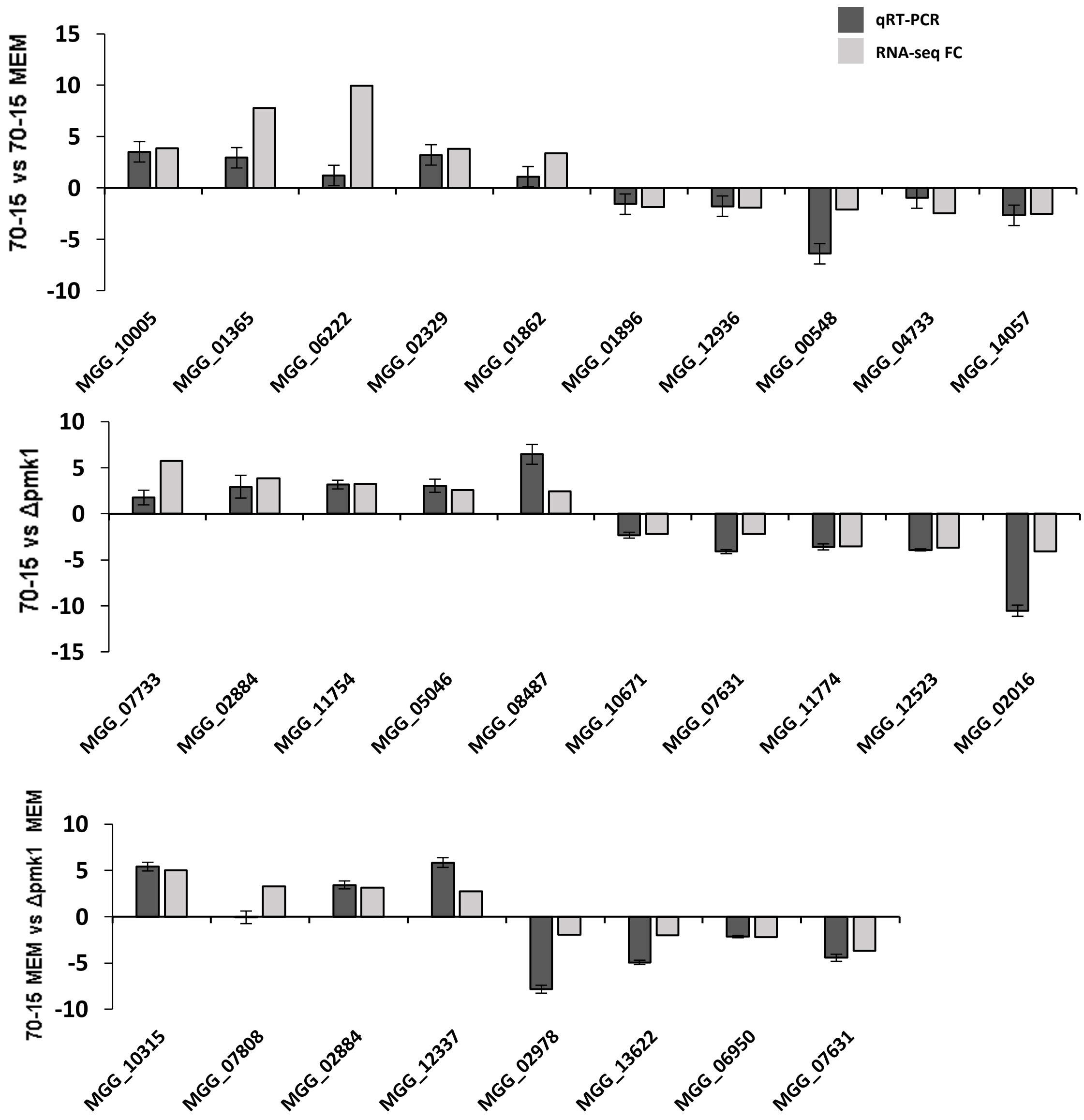


**Supplementary Fig. 7**. Validation of RNA-seq results by qRT-PCR for randomly selected DEGs. With a few exceptions, qRT-PCR results show positive correlation with RNA-seq data.

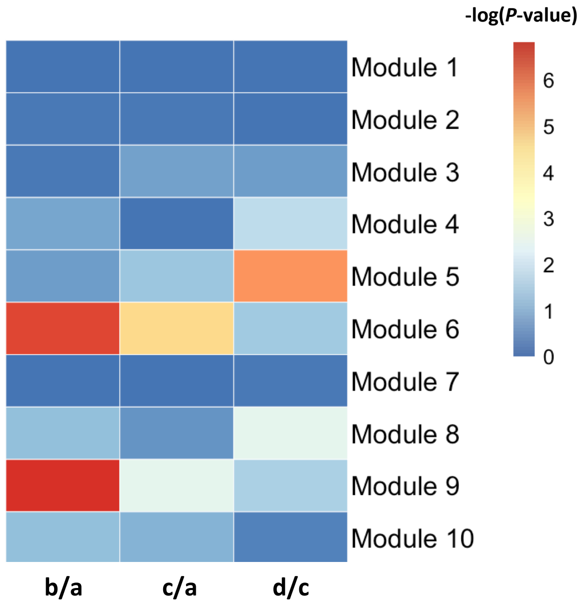


**Supplementary Fig. 8**. Heatmap showing significant enrichment of DEGs among ten expression modules defined by Yan et al. 2023 (Plant Cell). The DEG sets from b/a, c/a, and d/c were taken and compared with genes in individual modules for statistically significant enrichment. P-value was calculated using hypergeometric test.

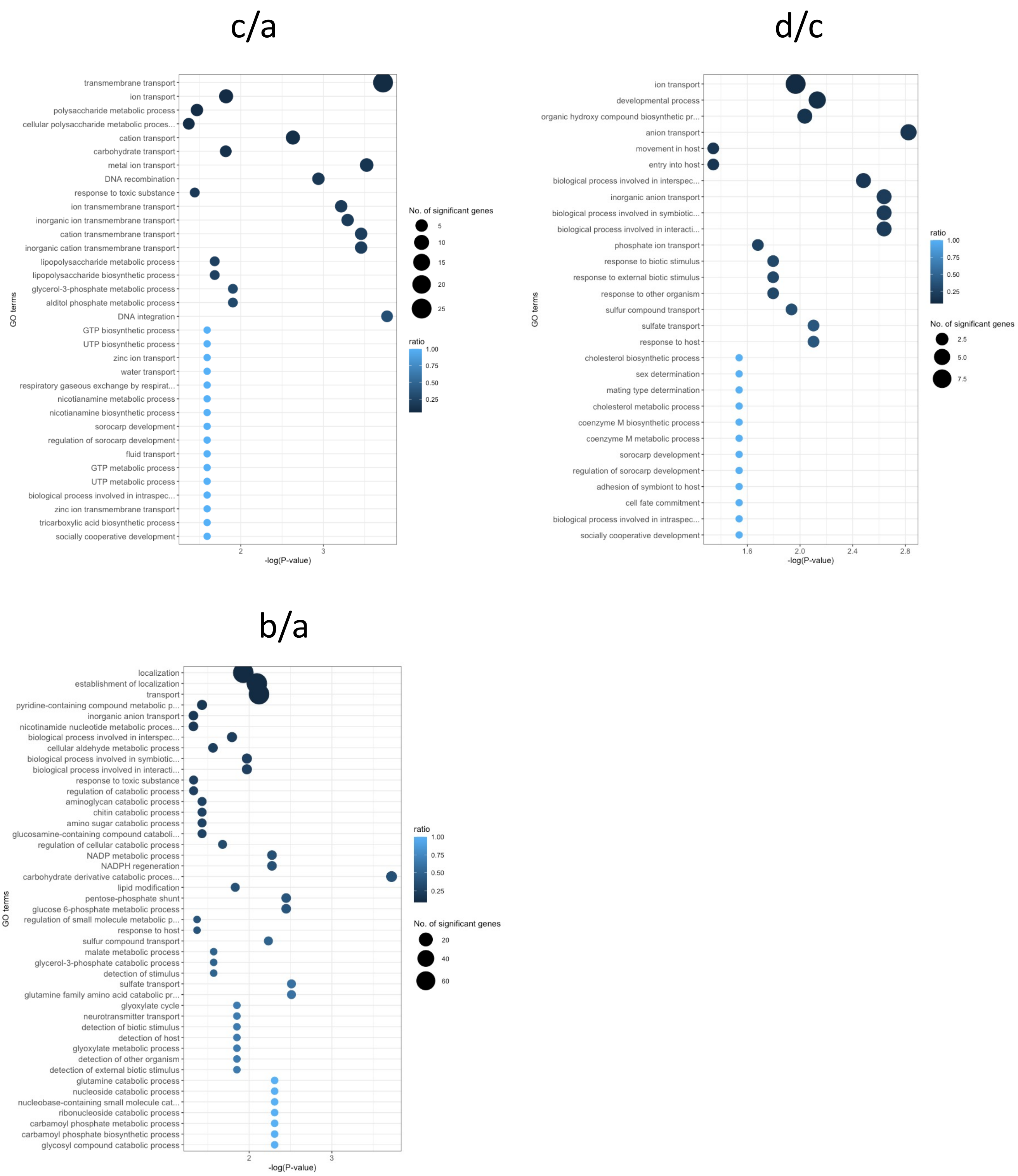


**Supplementary Fig. 9**. Summary of GO enrichment analysis (biological process) for selected terms. GO terms that are enriched with DEGs from c/a, d/c and b/a are shown (Fisher's exact test, P < 0.01). The area of the circle represents the number of genes assigned to the particular GO term. The color of the circle indicates the proportion of genes assigned to the GO term in our dataset among the total number of genes having that GO term in the genome.

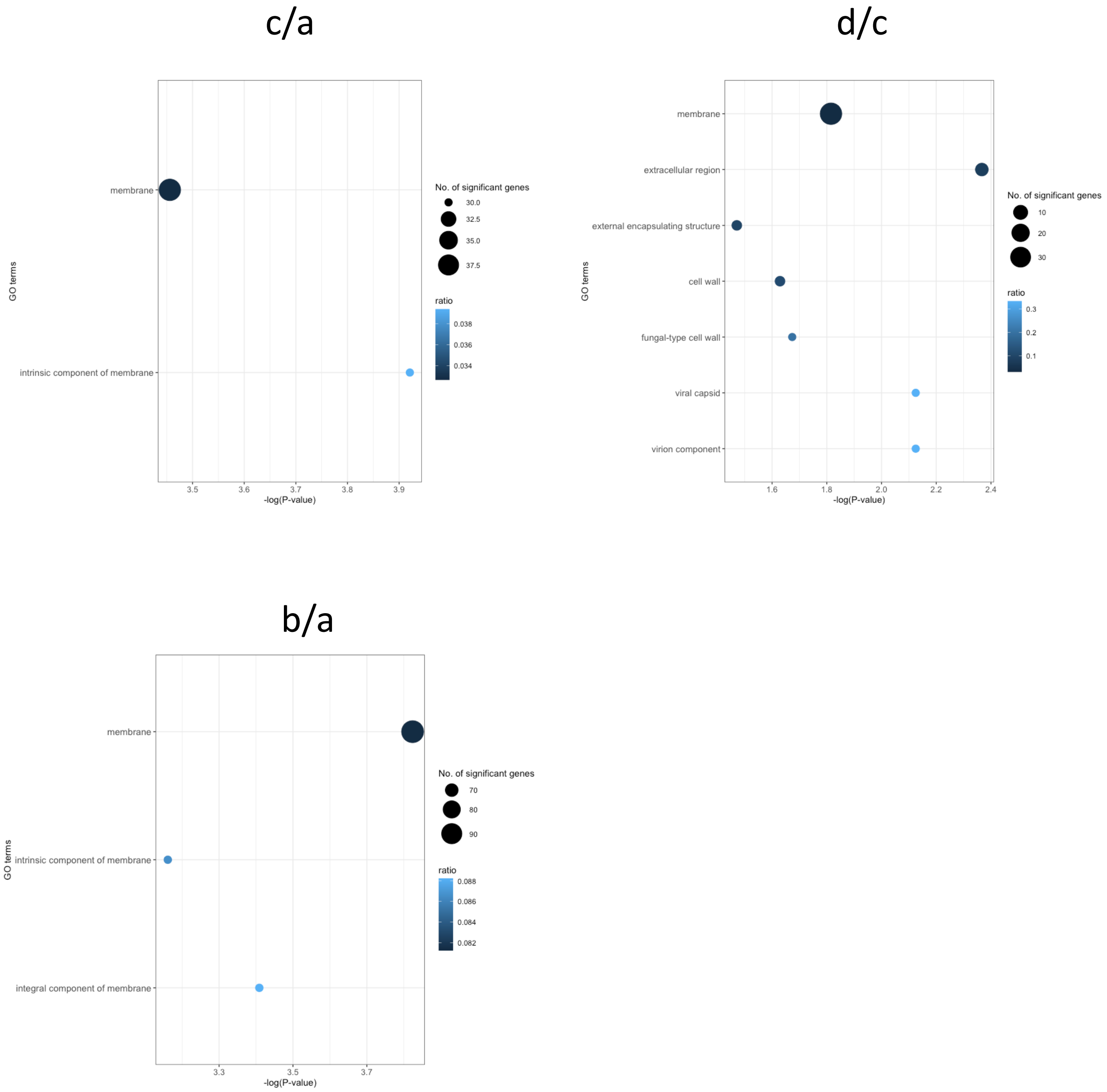


**Supplementary Fig. 10**. Summary of GO enrichment analysis (cellular component). GO terms that are enriched with DEGs from c/a, d/c and b/a are shown (Fisher's exact test, P < 0.01). The area of the circle represents the number of genes assigned to the particular GO term. The color of the circle indicates the proportion of genes assigned to the GO term in our dataset among the total number of genes having that GO term in the genome.

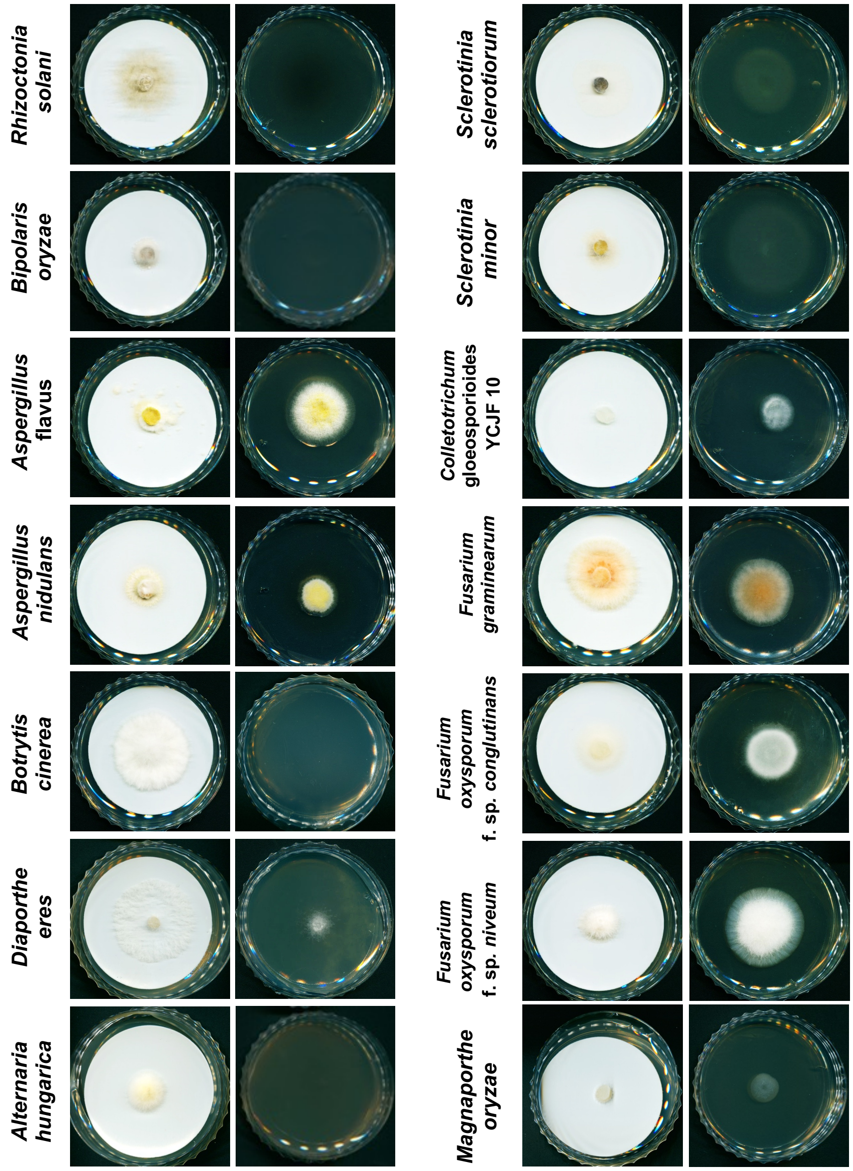


**Supplementary Fig. 11**. Assays for fungal species with different lifestyles. Saprobes, hemibiotrophs and necrotrophs were taken and tested for their ability to pass through the nitrocellulose membrane. Initial incubation period was adjusted according to the growth rate of each species.
